## Supplementary figures and images for "Enkephalin-mediated modulation of basal somatic sensitivity by regulatory T cells in mice"

### supplemental figures

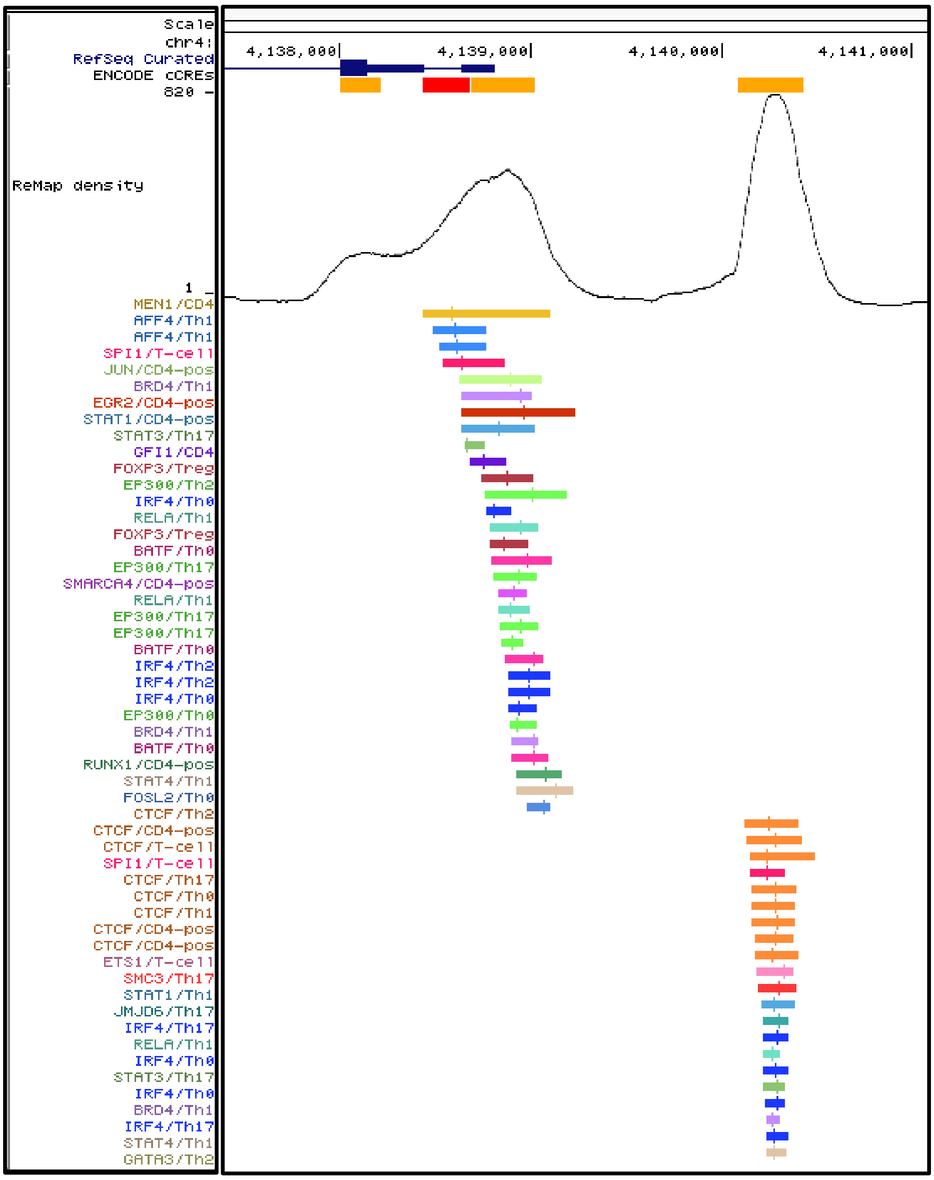

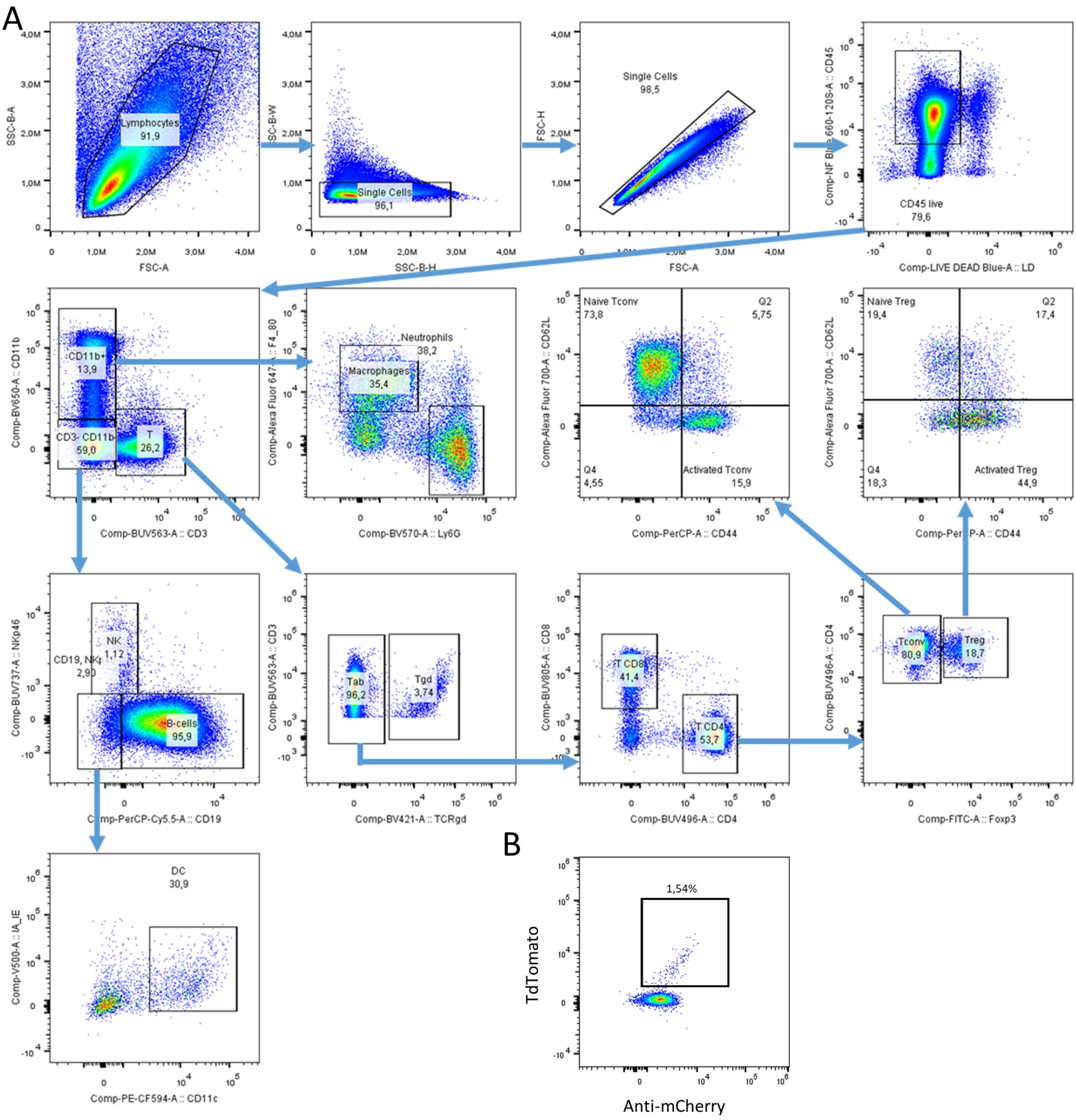

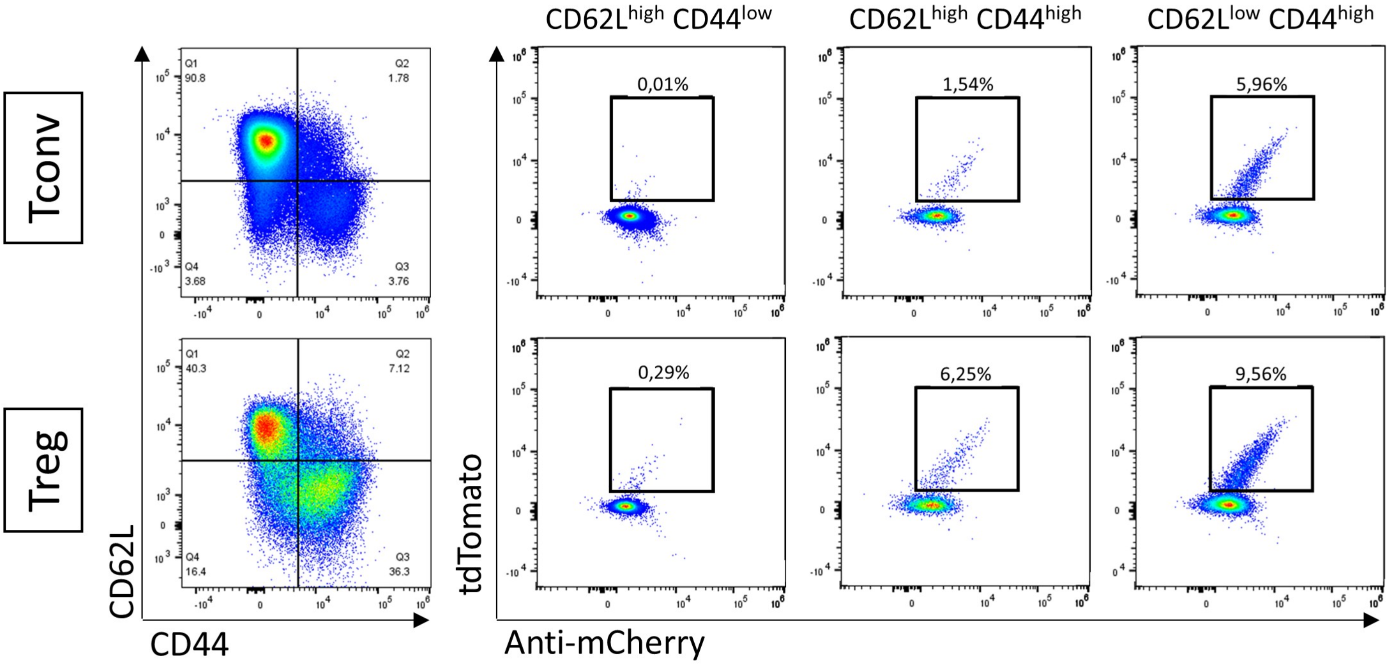

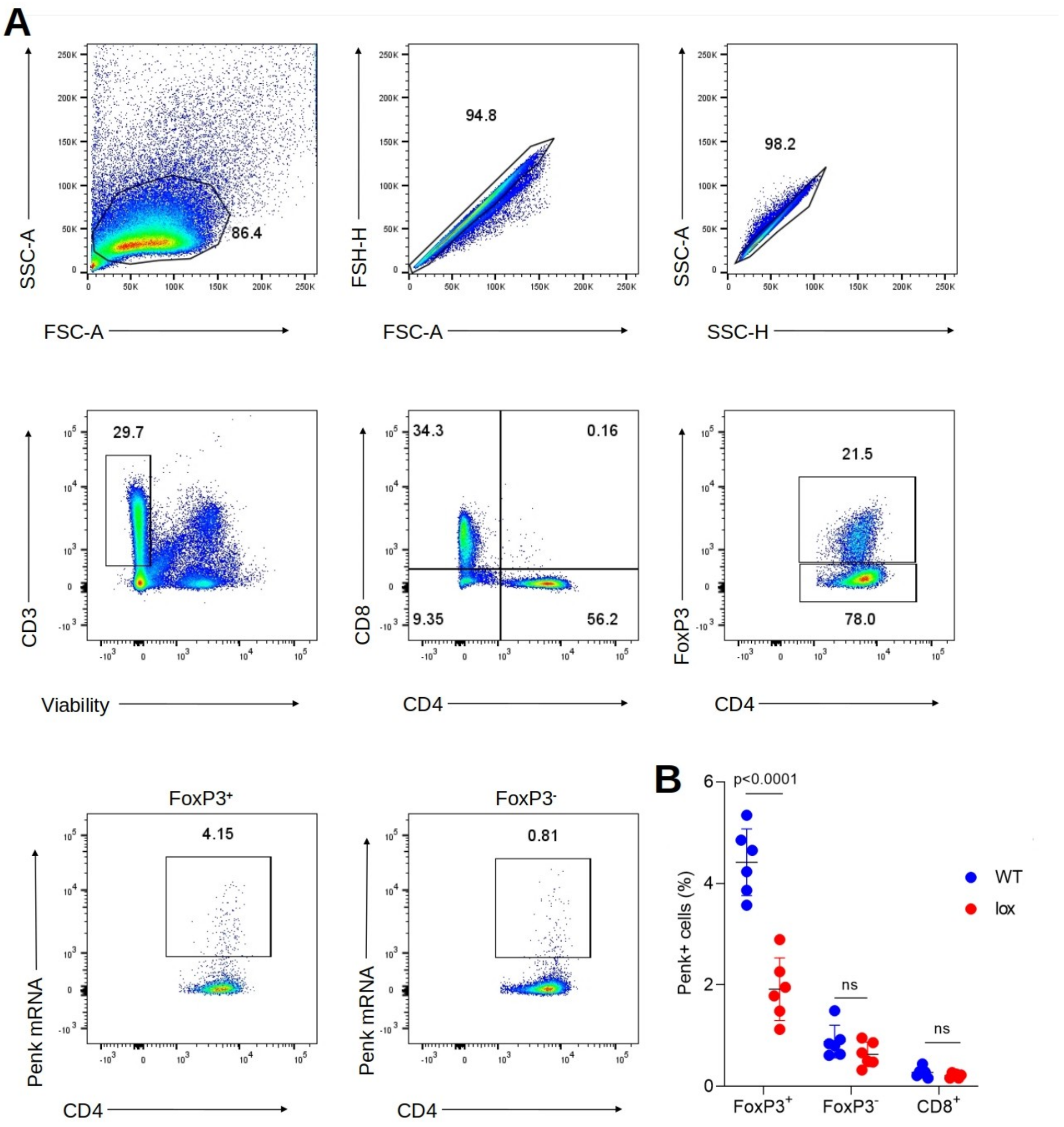

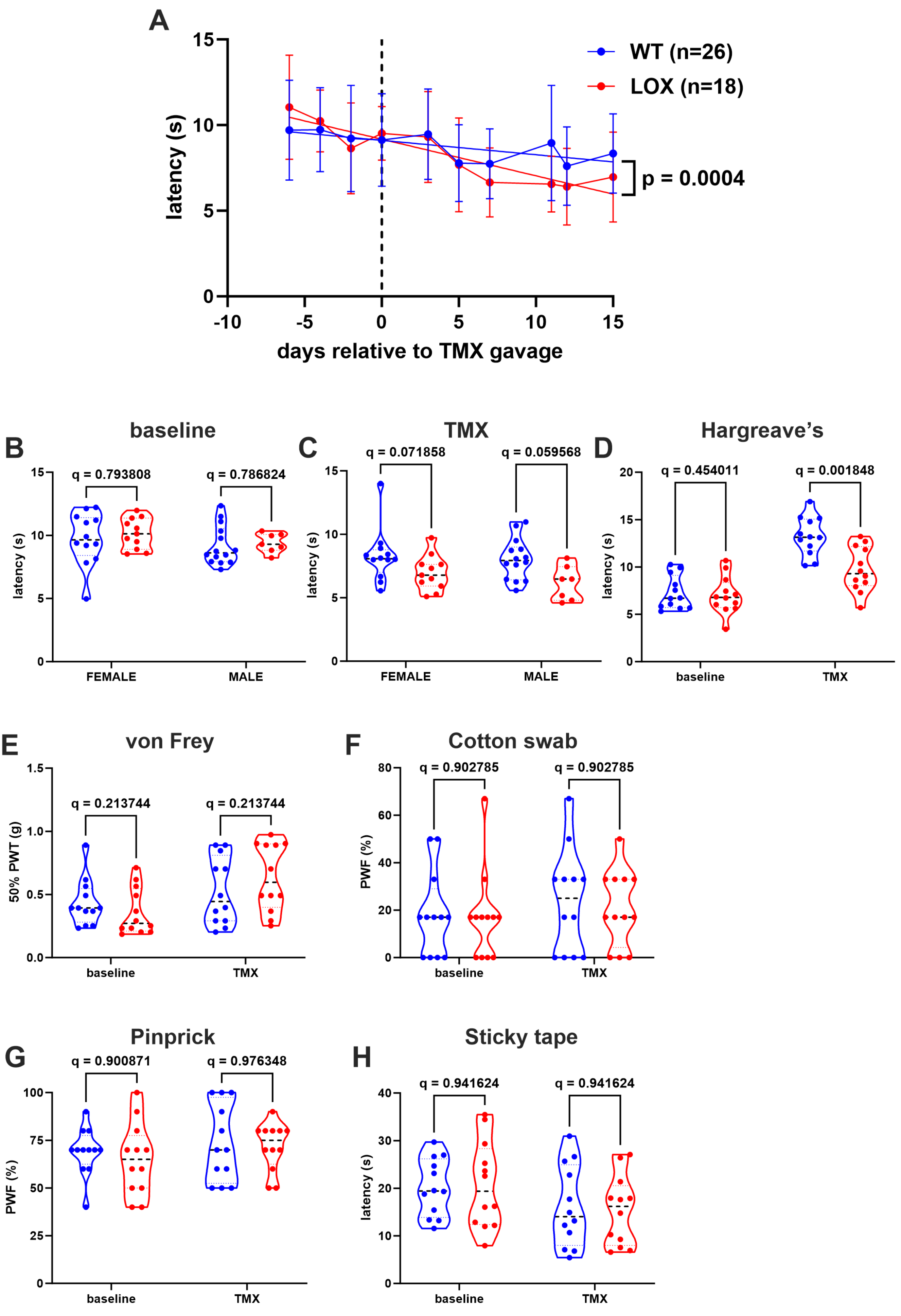
