## Supplementary material for "Enkephalin-mediated modulation of basal somatic sensitivity by regulatory T cells in mice": Table S2

Table S2 : Monoclonal antibodies used in the study of $Penk^{Cre}$ x $ROSA26^{tdTomato}$

| mAb or reagent (dye) | Provider | Clone | Dilution |
|---|---|---|---|
| Anti-mouse F4/80 (AF647) | Biolegend | BM8 | 1/400 |
| Anti-mouse CD62L (AF700) | BD Biosciences | MEL-14 | 1/400 |
| Anti-mouse CD25 (APC Fire 750) | Biolegend | PC61 | 1/100 |
| Anti-mouse CD4 (BUV 496) | BD Biosciences | GK1.5 | 1/400 |
| Anti-mouse CD3 (BUV563) | BD Biosciences | 145-2C11 | 1/400 |
| Anti-mouse CD69 (BUV661) | BD Biosciences | H1.2F3 | 1/200 |
| Anti-mouse Nkp46 (BUV737) | BD Biosciences | 29A1.4 | 1/200 |
| Anti-mouse CD8a (BUV805) | BD Biosciences | 53-6.7 | 1/800 |
| Anti-mouse Ly6G (BV570) | Biolegend | 1A8 | 1/200 |
| Anti-mouse CD11b (BV650) | Biolegend | M1/70 | 1/1000 |
| Anti-mouse CD45 (NovaFluor Blue 660-120S) | ThermoFisher | 30-F11 | 1/200 |
| Anti-mouse CD11c (PE CF594) | BD Biosciences | HL3 | 1/150 |
| Anti-mouse Ly6C (PE Cy7) | BD Biosciences | AL-21 | 1/1600 |
| Anti-mouse CD44 (PerCP) | Biolegend | IM7 | 1/200 |
| Anti-mouse CD19 (PerCP Cy5.5) | BD Biosciences | 1D3 | 1/400 |
| Anti-mouse IA/IE (V500) | BD Biosciences | M5/114.15.2 | 1/300 |
| Anti-mouse TCRγδ (BV421) | Biolegend | GL3 | 1/200 |
| Anti-mouse Ki67 (AF532) | ThermoFisher | SolA15 | 1/1600 |
| Anti-mouse mCherry (Pacific Blue) | ThermoFisher | 16D7 | 1/50 |
| Anti-mouse Foxp3 (FITC) | ThermoFisher | FJK-16s | 1/800 |
| Fixable viability dye (Live Dead Blue) | ThermoFisher | | 1/2000 |
